# Proof of concept study to develop a novel connectivity-based electric-field modelling approach for individualized targeting of transcranial magnetic stimulation treatment

**DOI:** 10.1101/2020.12.06.408856

**Authors:** Nicholas L Balderston, Joanne C Beer, Darsol Seok, Walid Makhoul, Zhi-De Deng, Tommaso Girelli, Marta Teferi, Nathan Smyk, Marc Jaskir, Desmond J Oathes, Yvette I Sheline

## Abstract

**Background:** Resting state functional connectivity (rsFC) offers promise for individualizing stimulation targets for transcranial magnetic stimulation (TMS) treatments. However current targeting approaches do not account for non-focal TMS effects or large-scale connectivity patterns. To overcome these limitations, we propose a novel targeting optimization approach that combines whole-brain rsFC and electric-field (e-field) modelling to identify single-subject, symptom-specific TMS targets.

**Methods:** In this proof of concept study, we recruited 91 anxious misery (AM) patients and 25 controls. We measured depression symptoms (MADRS/HAMD) and recorded rsFC. We used a PCA regression to predict symptoms from rsFC and estimate the parameter vector, for input into our e-field augmented model. We modeled 17 left dlPFC and 7 M1 sites using 24 equally spaced coil orientations. We computed single-subject predicted ΔMADRS/HAMD scores for each site/orientation using the e-field augmented model, which comprises a linear combination of the following elementwise products 1) the estimated connectivity/symptom coefficients, 2) a vectorized e-field model for site/orientation, 3) the pre-treatment rsFC matrix, scaled by a proportionality constant.

**Results:** In AM patients, our pre-stimulation connectivity-based model predicted a significant decrease depression for sites near BA46, but not M1 for coil orientations perpendicular to the cortical gyrus. In control subjects, no site/orientation combination showed a significant predicted change.

**Discussion:** These results corroborate previous work suggesting the efficacy of left dlPFC stimulation for depression treatment, and predict better outcomes with individualized targeting. They also suggest that our novel connectivity-based e-field modelling approach may effectively identify potential TMS treatment responders and individualize TMS targeting to maximize the therapeutic impact.

## Introduction

Major depressive disorder (MDD) is characterized by depressed mood or loss of interest or pleasure (1). Symptoms include feelings of worthlessness; indecisiveness or difficulty concentrating, change in weight, appetite, and sleep; psychomotor agitation; and suicidal ideation (1). According to the World Health Organization (WHO) World Mental Health (WMH) Surveys Initiative (2), lifetime prevalence rates within the United States average 16.9%, with women showing higher prevalence than men (20.2% vs. 13.2%) (2). The most common treatments for MDD include antidepressant medication, psychotherapy/cognitive behavioral therapy (CBT), or a combination of the two (https://www.nimh.nih.gov/health/statistics/major-depression.shtml). Although antidepressant use is prevalent, there are reports of mixed efficacy (3). Repetitive transcranial magnetic stimulation (rTMS) to the left dorsolateral prefrontal cortex (dlPFC) has been cleared by the US Food and Drug Administration (FDA), and has been shown to be effective as an antidepressant treatment in treatment resistant depression, a patient population not responsive to or tolerant of previous antidepressant medication trials (4–6). Although many studies find a significant reduction in depressive symptoms following rTMS treatment (7–13), there have also been studies finding limited efficacy for major depression (14, 15), leaving room for improvement.

One possible explanation for the heterogeneity in TMS treatment response for major depression is that the standard scalp-based targeting approach (i.e. the 5-cm rule) does not account for individual variations in cortical anatomy, and often selects regions outside the dlPFC, which may explain some of the modest therapeutic effects (16). Image-guided TMS incorporating MRI-neuronavigation, taking into account individual differences in brain anatomy, can target specific functional brain networks with greater accuracy (17, 18). Using structural MRI to locate a specific site at the junction of BA 9 and 46 has also shown greater clinical efficacy in reducing depression than the traditional 5-cm rule (15). More recent approaches have used functional MRI (fMRI) to target regions that are downstream of the dlPFC, like the subgenual anterior cingulate cortex (sgACC) (19–21) that are thought to be dysfunctional in MDD patients (22–25). Key to this approach is to use functional connectivity of the downstream site to identify the dlPFC site with the strongest connectivity. Finally, using e-field modelling can improve accuracy even further. Indeed, a recent report estimates the difference between conventional neuronavigated targeting and e-field informed target to be up to 14 mm (27).

Connectivity-based approaches represent the current cutting-edge in the field and are an improvement on the standard scalp-based targeting approach used clinically. However as currently implemented, they typically suffer from two fundamental limitations, which we address in the current work using a novel computational model based on whole brain functional connectivity and electric-field modelling. The first limitation is that by focusing on a single downstream region to define the TMS target they do not account for off-target effects in other downstream regions. To address this limitation, we use electric-field modelling to estimate the changes in connection strength across the entire brain, thus accounting for changes in connectivity that would otherwise be considered “off-target”. The second limitation is that most connectivity-based targeting approaches do not typically quantify the link between functional connectivity and symptoms, making it difficult to make a quantitative prediction about symptom change following neuromodulation. To address the second limitation, we use a PCA regression approach to quantify the relationship between functional connectivity and depression symptoms, and use the resulting coefficients to estimate the net effect of TMS on connectivity and thus symptoms. The overall concept is to generate a predicted symptom change score following neuromodulatory TMS at a particular site and stimulation. We then iterate the model across sites and orientations to find the site/orientation combination that leads to the maximal reduction in MDD symptoms. Importantly, because this approach is data-driven it has an added benefit of being both site and symptom independent.

The current work is a proof of concept demonstration of this novel targeting approach using structural and functional MRI collected from the Dimensional Connectomics of Anxious Misery dataset (See Seok et al., 2020 (26) for full characterization of cohort). The goal is to characterize the relationship between symptoms and connectivity, then model the effect of TMS on connectivity in order to predict the change in symptoms that would be likely to occur given stimulation at a particular site and orientation. In the current work, we apply this modeling approach to two targets, a positive control left dlPFC and a negative control motor cortex (M1), based on targeting guidelines from the literature (See Supplemental Tables 1 and 2). We use bootstrapping to estimate the reliability of the targets identified using this approach at the single-subject level, and permutation testing to determine whether this approach makes group-level predictions that are consistent with the literature (28–30).

**Table 1:**
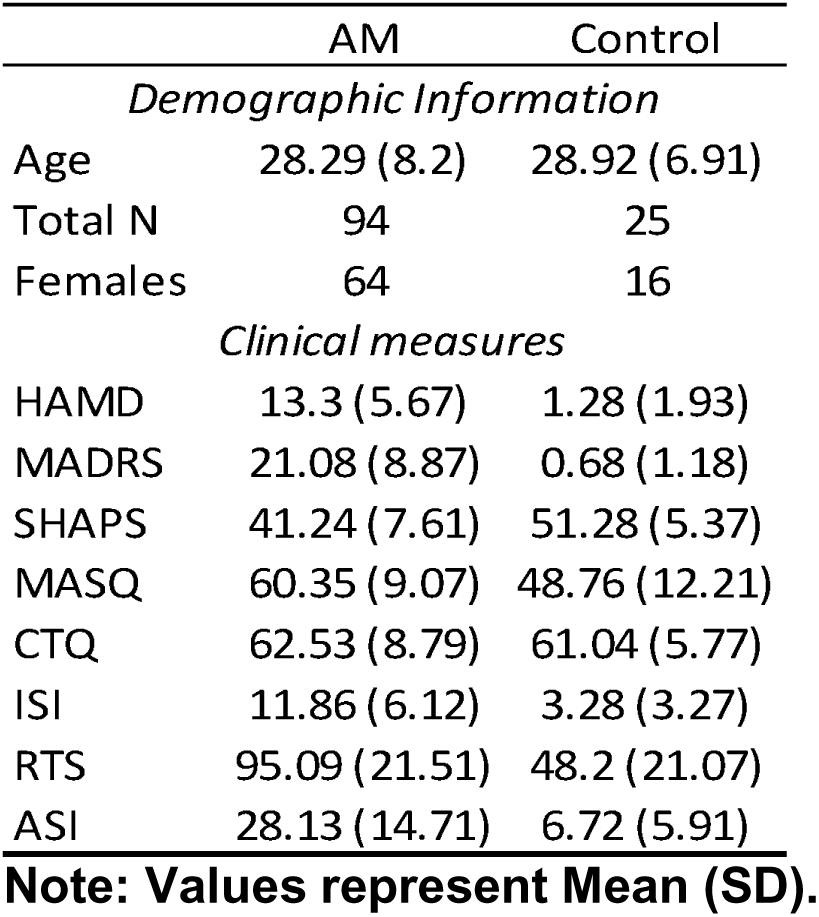
Demographic information and clinical measures

|  | AM | Control |
| --- | --- | --- |
| *Demographic Information* |  |  |
| Age | 28.29 (8.2) | 28.92 (6.91) |
| Total N | 94 | 25 |
| Females | 64 | 16 |
| *Clinical measures* |  |  |
| HAMD | 13.3 (5.67) | 1.28 (1.93) |
| MADRS | 21.08 (8.87) | 0.68 (1.18) |
| SHAPS | 41.24 (7.61) | 51.28 (5.37) |
| MASQ | 60.35 (9.07) | 48.76 (12.21) |
| CTQ | 62.53 (8.79) | 61.04 (5.77) |
| ISI | 11.86 (6.12) | 3.28 (3.27) |
| RTS | 95.09 (21.51) | 48.2 (21.07) |
| ASI | 28.13 (14.71) | 6.72 (5.91) |

**Note:** Values represent Mean (SD).

## Materials and Methods

### Participants

One hundred eighteen participant (Age (years): M=28.42, SD=7.92) entries were pulled from the Dimensional Connectomics of Anxious Misery dataset (26), and included anxious misery participants (93 total, 64 were females) and healthy controls (25 total, 16 were females; See Table 1). Mean (SD) age of the anxious misery cohort was M=28.29 (8.2) and of healthy controls M=28.92 (6.91). Patients included in the anxious misery cohort were selected on the basis of their score on the NEO Five-Factor Inventory (NEO FFI) (31) rather than on diagnosis in keeping with the RDoC concept of finding symptom domains across diagnoses (32, 33). In post-hoc analyses, the cross-diagnostic anxious misery cohort included a range of different primary diagnoses, namely major depressive disorder (N = 41), persistent depressive disorder (N = 9), social anxiety disorder (N = 5), generalized anxiety (N = 21), and PTSD (N = 17).

Basic Inclusion/Exclusion criteria included MRI contraindications, histories of certain neurological or cognitive disorders and events, histories of exclusionary psychiatric conditions or any other factors that may have affected patient safety (See Seok et al. (26) for complete list). In addition, participants in the anxious misery cohort needed to score a minimum of one (1) standard deviation above a point estimate of the general adult population mean (distributions for males and females were different) in the NEO FFI (31) in order to be included in the study (26). Conversely, healthy controls needed to score one (1) standard deviation or more below the mean point estimates to be eligible to participate. Subjects were recruited by means of IRB-approved advertisement and phone calls. All participants signed an informed consent form, and the protocol was approved by the Institutional Review Board for human subject research at the University of Pennsylvania.

### Procedure

On the initial visit, written consent was obtained from all eligible participants prior to enrolling in the study, followed by a baseline visit with study staff to ascertain their medical history and demographic information. In addition, a research version of the Structured Clinical Interview for DSM-5 (SCID-5-RV) (1) was conducted by a trained member of the staff to document their psychiatric diagnosis and history, and the remaining assessments (See Below) were conducted. In a follow-up visit, a 2-hour fMRI scan was conducted, where structural, diffusion, resting state, and task-based fMRI data were collected as part of the larger project.

### Assessments

Many clinical measures were used to identify the behavioral and cognitive features of anxious misery. In this project, eligible participants completed 17 self-report and 3 clinician-administered measures. The clinician-administered measures included the Montgomery-Åsberg Depression Rating Scale (MADRS) (34) and the Hamilton Depression Rating Scale (HAMD) (35) in order to assess depression severity.

### MRI Scans

MRI data was acquired as part of the Dimensional Connectomes of Anxious Misery project (26), one of the Connectomes Related to Human Diseases (CRHD) studies (https://www.humanconnectome.org/disease-studies). Participants were scanned on a Siemens Prisma 3T using a 64-channel head coil. Structural T1-weighted images were acquired using a magnetization-prepared rapid acquisition with gradient echo (MPRAGE) sequence with TR=2400ms, TE=2.22ms and flip angle of 8 degrees. 208 slices were acquired with a voxel resolution of 0.8mm isometric, resulting in an FOV of 256 x 240 x 167mm. T2-weighted images were acquired using a variable-flip-angle turbo-spin echo (TSE) sequence with TR=3200ms and TE=563ms, with the same voxel resolution and FOV as the T1w acquisition. Resting state fMRI data were acquired with a multi-band acceleration of 8, TR=800ms, TE=37ms and flip angle of 52 degrees. Whole-brain coverage was achieved with 72 slices and a voxel resolution of 2.0mm isometric, resulting in an FOV of 208 x 208 x 144mm. Each resting state scan was paired with another run with the opposite phase encoding direction (AP-PA), and two such pairs were acquired, resulting in 22:24 minutes of data (5:46 min x 4 runs) for each participant. Spin echo field maps were also acquired in opposite phase encoding directions in order to correct susceptibility distortions. During the resting state scan, participants were shown a white screen with a black crosshair in the center and were instructed to remain still with their eyes open and to blink normally.

### MRI preprocessing

#### Computational head modelling for e-field calculations

We used the SimNIBS (Version 2.1) software package to generate 3D head and coil geometries using the finite element method (FEM) (36). Individualized head models were created using T1 and T2 structural MRIs that were segmented into scalp (mri2mesh), skull, CSF, gray matter, and white matter volumes, then converted to tetrahedral meshes using a Gmsh subroutine packaged in SimNIBS (37).

#### E-field calculations

E-field models were conducted for each site/orientation combination (See Below), and E-norm (i.e. magnitude) component of the electric field at each surface node was used to model the potential TMS effects (38).

### fMRI preprocessing

Data were preprocessed using the Human Connectome Project Pipelines v4.0.1 (retrieved from https://github.com/Washington-University/HCPpipelines/releases/tag/v4.0.1). A full description of preprocessing steps is provided elsewhere (39). In brief, anatomical preprocessing steps included gradient nonlinearity distortion correction, co-registration of T1w and T2w images, bias-field correction using spin-echo field maps and spatial normalization to the Montreal Neurological Institute (MNI) template. Functional image preprocessing steps included removal of spatial distortions via gradient nonlinearity corrections, correction of participant motion through volume realignment, susceptibility distortion correction using dual-phase encoded spin-echo corrections, registration of fMRI data to T1 space, subsequent transformation to MNI-space and removal of extra-parenchymal voxels.

Timeseries analyses using volumetric data were further conducted using the eXtensible Connectivity Pipeline (XCP Engine) (40). The workflow is summarized as follows: (i) removal of the 10 initial volumes (8 seconds) to achieve signal stabilization, (ii) demeaning and removal of quadratic trends using a general linear model to account for scanner drift, (iii) intensity despiking using 3dDespike from AFNI (41), (iv) bandpass temporal filtering of time series between 0.01 Hz and 0.08Hz using a first-order Butterworth filter (42), (v) regression of nine confounding signals (six motion parameters + global signal + mean white matter signal + mean cerebral spinal fluid signal) and as well as the temporal derivative, quadratic term and temporal derivatives of each quadratic term (resulting in 36 regressors total) (43), and (vi) spatial smoothing with SUSAN from FSL (44) using a 6mm FWHM kernel. Voxelwise timeseries were then downsampled to the 333 parcels in the Gordon atlas (45) and pairwise interparcel connectivity was calculated using Z-transformed Pearson correlations.

### Model steps

#### PCA regression

For the first step in our analysis (See Figure 1 and Appendix for detailed equations), we used principal components analysis (PCA) regression to model the relationship between functional connectivity and symptoms (37, 38, Eq. A2). We began by conducting a principal components analysis on the functional connectivity data for each group (i.e. AM and control) to reduce noise and minimize collinearity (48). We restricted attention to the subset of components that explained largest proportions of variability, with the assumption that these contain most of the relevant signal. For this we used a geometric approach that identified the eigenvalue at the elbow of the scree plot (i.e. the eigenvalue furthest from the hypotenuse connecting the first and last eigenvalues). The principal component scores for each of these components were then used to model symptom scores (MADRS and HAMD) in a multiple linear regression. The resulting coefficient estimates were then projected back into feature space (Eq. A3). The result was a vector where each element represents the coefficient for the corresponding functional connection in a linear model predicting symptom score.

**Figure 1.**
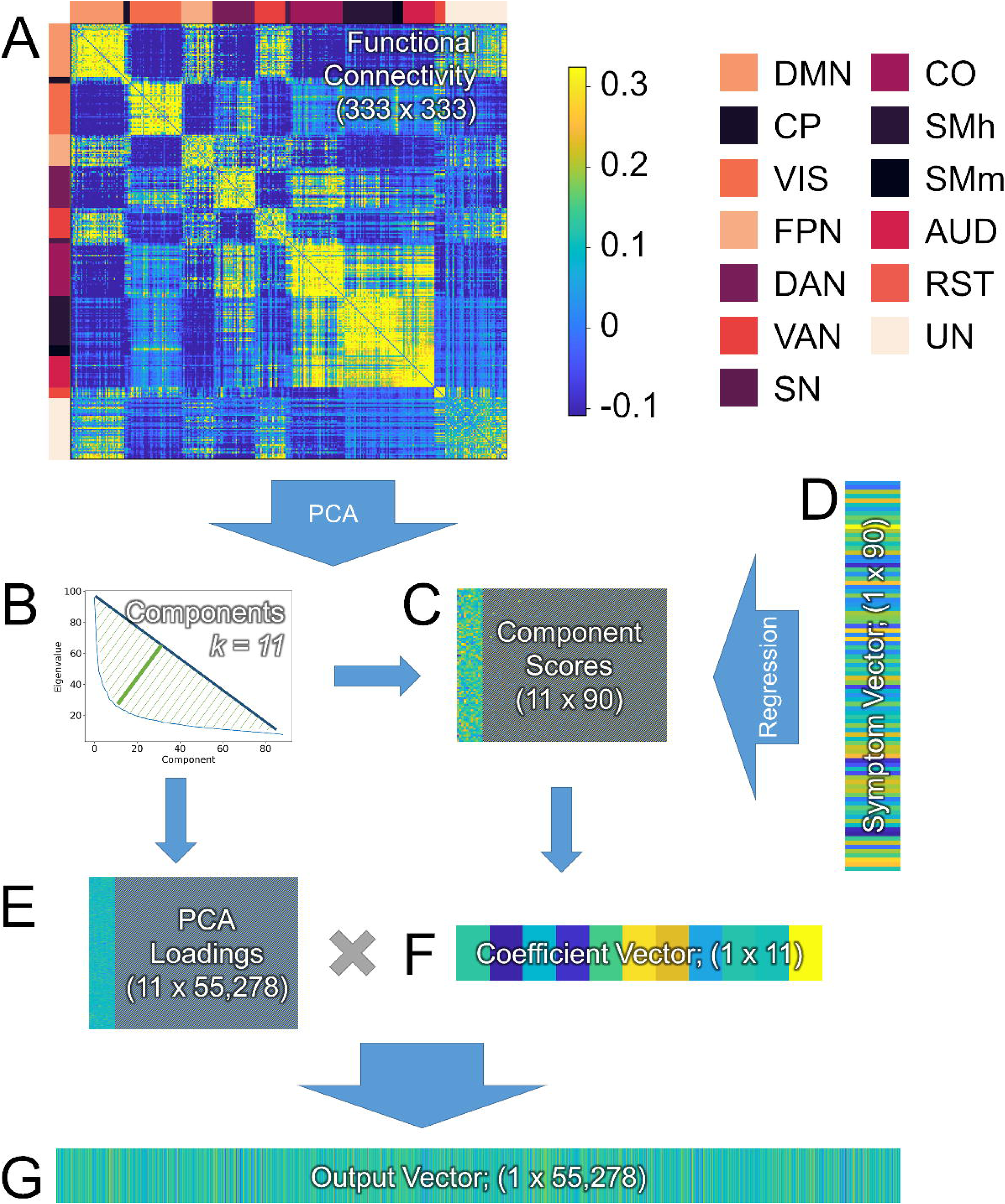
The PCA-Regression data reduction approach used to summarize the relationship between symptoms and connectivity in the current model. **A**) Resting state functional connectivity (rsFC) is calculated using Pearson’s correlation across all subjects for all regions in the Gordon atlas (45). **B**) A principal component analysis (PCA) is used to identify orthogonal components in the rsFC data, and a geometric approach is used to identify a minimal number of components that explain a maximal proportion of the variability. **C**) Component scores for the selected components are extracted and entered into a multiple linear regression to predict symptoms (**D**). The PCA loadings from the selected components (**E**) are combined with the coefficient vector from the regression (**F**) using matrix multiplication to create the output vector (**G**), which is used to represent multiple regression coefficients projected into the rsFC feature space. **Network Color key: DMN =** Default Mode Network; **CP =** CinguloParietal; **VIS =** Visual; **FPN =** FrontoParietal Network; **DAN =** Dorsal Attention Network; **VAN =** Ventral Attention Network; **SN =** Salience Network; **CO =** CinguloOpercular; **SMh =** SomatoMotor (hand); **SMm =** SomatoMotor (mouth); **AUD =** Auditory; **RST =** RetrosplenialTemporal; **UN =** Unassigned nodes.

#### Selection of sites and orientations

To characterize the dlPFC, we defined a series of 17 sites along the anterior-to-posterior gradient of the L dlPFC, which was prioritized based on prior clinical efficacy findings (See Supplemental Table 1). This site vector was anchored by 3 therapeutic sites described in (30) ([-41, 16, 54] based on 5cm rule, [-36, 39, 43] BA9, [-44, 40, 29] BA46), and extends 12 mm posterior to the 5mm site and 12 mm anterior to the BA46 site. To characterize the hand knob, which was used as a negative control region, we defined a series of 7 sites along the medial to lateral axis of the primary motor cortex in the left hemisphere (See Supplemental Table 2). This anchor was centered on the hand knob, which was identified from Neurosynth using an association test with the search term “hand”. The center of mass for the primary motor cortex cluster was used as the hand knob coordinate ([−36, −24, 58]). At each site, we generated 24 different e-field models. For each model, the roll and pitch of the TMS coil was defined perpendicular to the scalp. The yaw vector was selected from 24 possible orientation vectors spaced at 15-degree increments.

#### E-field connectivity matrices

For each participant, the e-field map was downsampled (i.e. averaged across voxels in each parcel) to the Gordon atlas (45) and concatenated into a 1 x 333 vector. This vector was then sorted by e-field magnitude to resemble a scree plot, and thresholded at the value furthest from the hypotenuse connecting the first and last value in the vector (See Figure 2). To estimate how stimulation at a specific site and orientation might affect connectivity, the downsampled e-field vector was used to form an a 333 x 333 matrix where the values in the matrix represent the average current induced in the ROIs for each connection.

**Figure 2.**
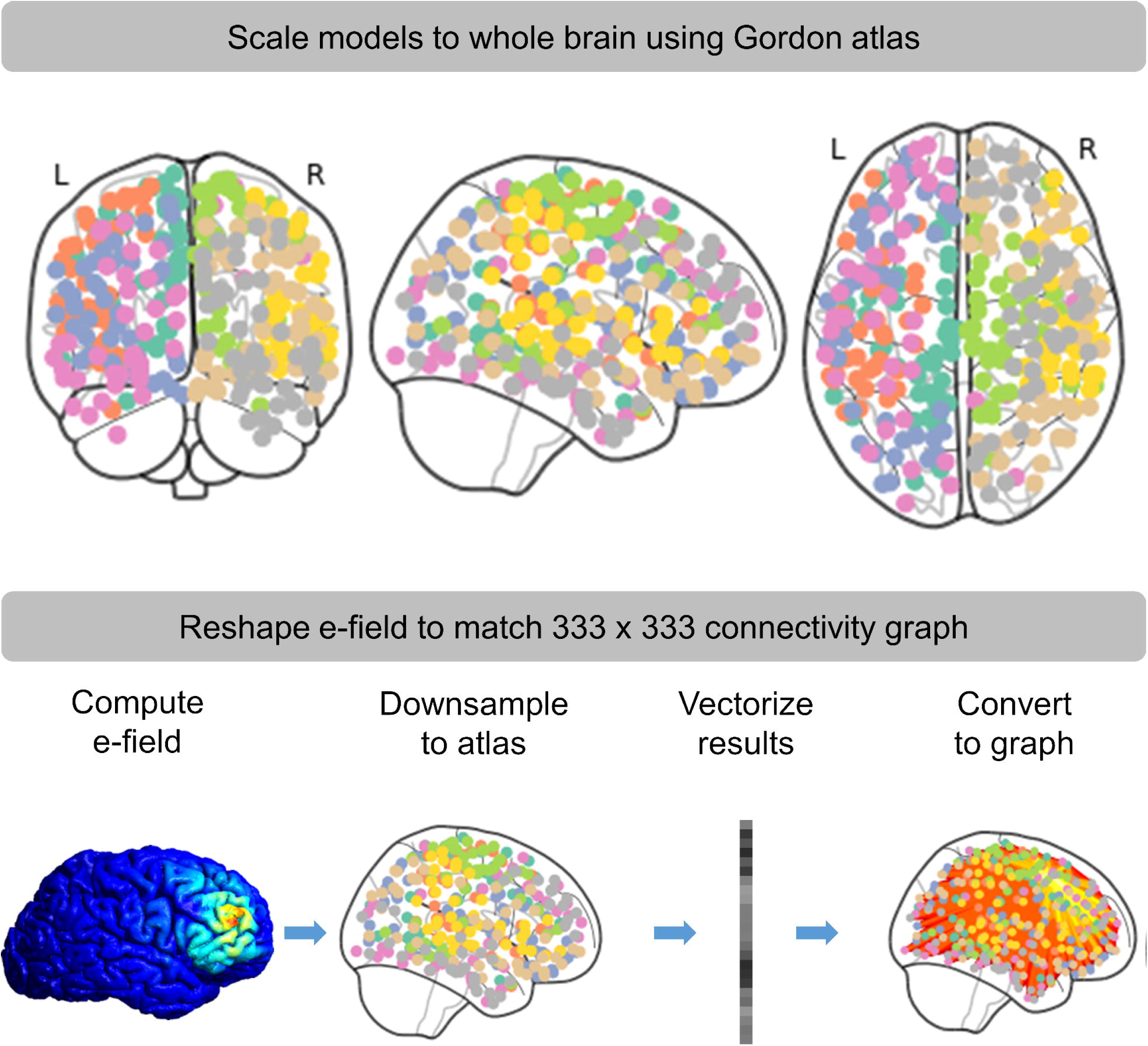
Methods used to summarize electric (e)-field models in connectivity space. Normalized e-field models were first downsampled to the Gordon atlas (45). The results were then converted to a 1 x 333 vector. This vector was then used to form an a 333 x 333 matrix where the values in the matrix represent the average current induced in the ROIs for each connection.

### E-field/connectivity symptom estimates

The predicted neuromodulatory effect of TMS on symptoms was modelled using each participant’s e-field model matrix (49), the single-subject connectivity matrix, and the estimated group-level coefficients derived from the symptom/connectivity regression, according to Equation A2-3. These calculations were iterated across all sites and orientations and spatially smoothed using a Gaussian kernel (SD = size(heatmap)/10; See MATLAB function smoothdata) to create the heatmap shown in Figure 3, where each value in the heatmap represents the predicted relative change in symptoms (sans application of an empirically-determined constant scaling factor *C*) for that subject given hypothetical TMS administered at that particular site and orientation.

**Figure 3.**
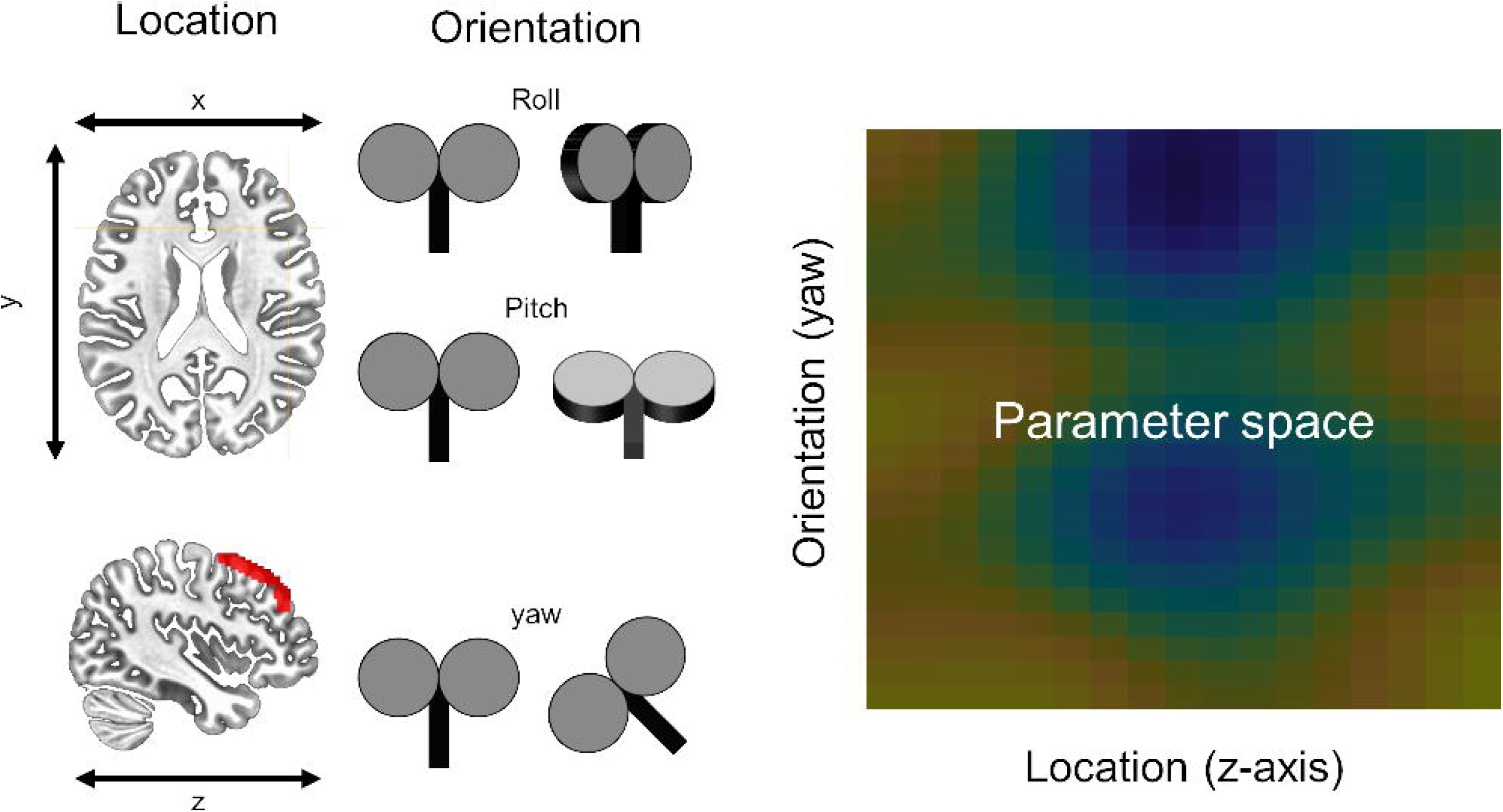
Iteration of model across site and orientation. To understand how placement and orientation of the TMS coil might impact symptoms, we computed our model across multiple sites and orientations. Sites were defined at equally spaced points along the anterior to posterior axis of the middle frontal gyrus. Roll and pitch were defined orthogonal to the scalp at each stimulation site. Multiple equally spaced yaw vectors were defined at each stimulation site, representing all possible coil orientations. Electric (e)-field models were conducted at each site/orientation combination, entered into our symptom prediction model, and the results were plotted in this site x orientation heatmap.

### Statistical analysis

#### Individual subject predictions

One goal of this project was to determine whether the modelling approach above could be used to make targeting decisions for the application of TMS to single subjects. Accordingly, to be effective our model would need to show a systematic effect of site and orientation on predicted symptom change in order to be used for single-subject targeting. To examine this possibility, we created single-subject heatmaps corresponding to the predicted relative change in symptoms as a function of site and orientation, and identified the point in these heatmaps that yielded the maximum predicted relative decrease in symptom scores, corresponding to the optimal site/orientation of stimulation for that subject.

We then used bootstrapping to estimate the reliability of this targeting approach. For each of the 10,000 iterations of the bootstrapping analysis, we sampled subjects with replacement from the original distribution, recalculated the PCA regression and the corresponding 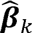 vector (See Appendix: EQ5). Next we recreated the individual subject heatmaps, and identified the optimal stimulation site. Across bootstrap iterations we calculated the Euclidean distance between the single-subject original optimal site/orientations and the bootstrap-calculated optimal site/orientations. Then for each subject, in order to form a confidence region, we identified the value corresponding to the 95^th^ percentile from the distance metric bootstrap distribution, which was used to display the reliability of the single-subject target in the heatmaps.

#### Group-level predictions

To determine whether our novel modelling approach predicted significant changes at the group level, for each site/orientation we conducted one-sample t-test of the predicted change in symptoms (i.e. predicted post treatment symptom score – baseline symptom score) against zero. A significant negative t-score would indicate a predicted decrease in symptoms at the group level. Accordingly, the null hypothesis for each site/orientation combination is that the model predicts no significant group-level change in symptoms following a hypothetical treatment at that site/orientation. We repeated this process for all site/orientation combinations, and generated heatmaps from the resulting t-scores. We used cluster-based permutation testing to correct for multiple comparisons across site and orientation (50). First we conducted 10,000 permutations where the sign of the predicted symptom change was randomly flipped across subjects and permutations. For each permutation, we calculated t-scores at each site/orientation combination within the heatmap. We thresholded these t-scores using a 2-tailed alpha of 0.025. Next we identified clusters of t-scores surviving the threshold, summed the values across elements within the cluster, identified the cluster with the maximum summed t-score, and built a null distribution with these max summed t-score values. We then identified the critical values corresponding to the 1.25^th^ and 98.75^th^ percentiles (with alpha=0.025 to correct for the number of regions of interest tested) of the max summed t-score distribution generated from the permutation procedure. Finally, a two-step thresholding process was implemented to threshold the group-level statistical maps. First, the individual t-scores were thresholded using an alpha of 0.025. Then, summed t-scores from the surviving clusters from these heatmaps were thresholded using the critical values from the max summed t-score null distribution.

## Results

### PCA regression

Separate PCA regression analyses were conducted for the anxious misery group and the control subjects. For the AM group, 11 signal components were selected, and these components explained 31% of the variability in the functional connectivity signal (See Supplemental Figure 1). For the control group, 6 components were selected, and these components explained 40% of the variability in the functional connectivity signal (See Supplemental Figure 2).

### Individual subject predictions

Single subject heatmaps from the AM group show clear systematic patterns of predicted symptom change as a function of both site and orientation (See Figure 4), suggesting that this modelling approach can be used to simultaneously optimize these two parameters at the single subject level. These patterns are characterized by punctate “cool spots” in the heatmap that identify a clear site/orientation combination that yields maximal predicted symptom reduction. To characterize the variability in our targeting approach, we conducted 10,000 bootstrapping iterations and identified the optimal site/orientation combination on each. We then calculated the Euclidean distance between the bootstrap-derived values and the original values and plotted a circle with an area that corresponds to the 95^th^ percentile in the bootstrapped Euclidean distance distribution on the single subject graphs shown in Figure 4 and Supplemental Figure 3. Importantly, this pattern seems to be specific to those in the AM group as control subjects do not show the characteristic “cool spots” in their heatmaps (See Supplemental Figure 3).

**Figure 4.**
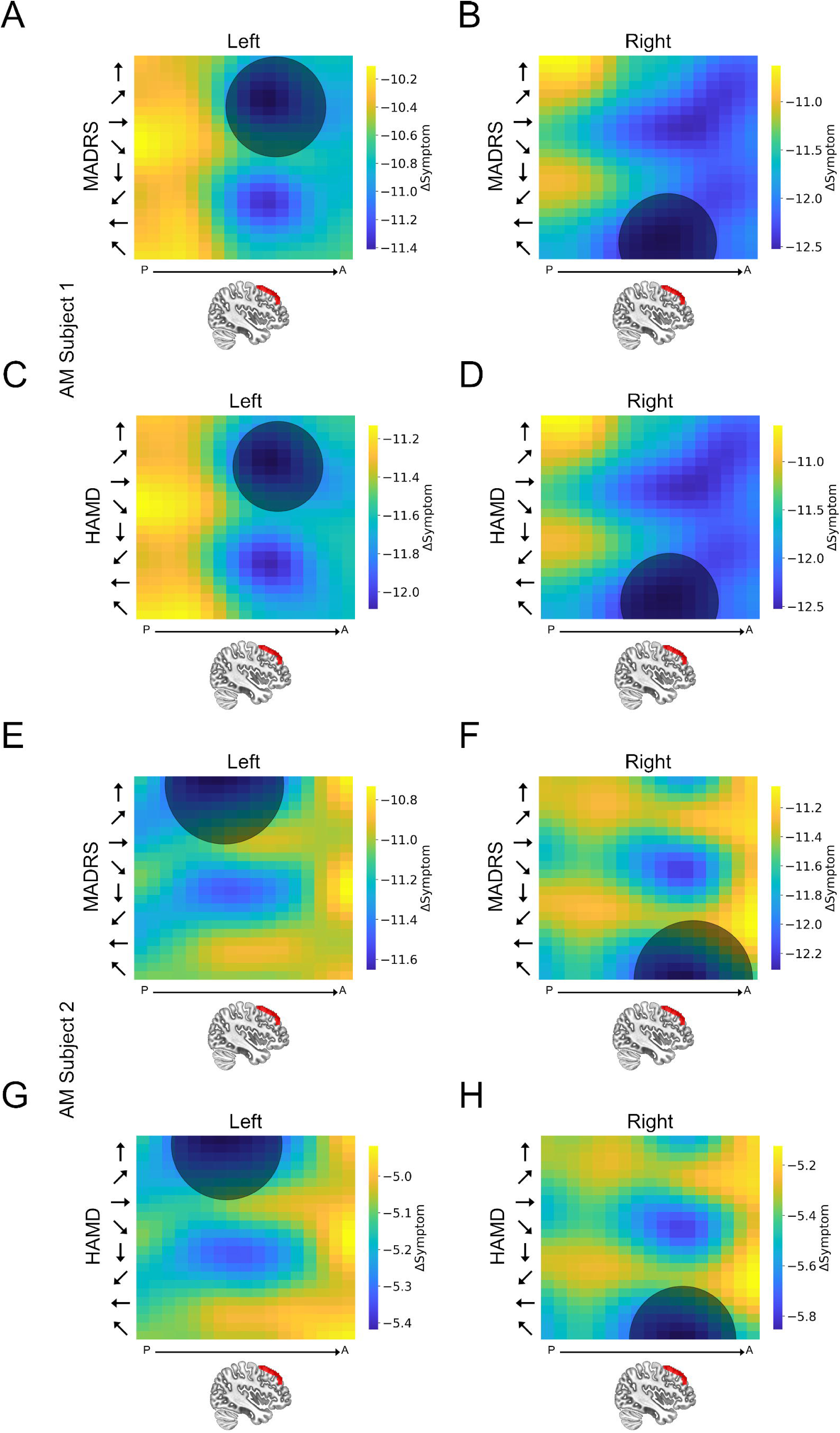
Individual subject heatmaps plotting dlPFC predictions for the anxious misery group. (**A, C, E, G**) Heatmaps representing the predicted MADRS and HAMD scores following a hypothetical course of TMS treatment to the left dlPFC. (**B, D, F, H**) Heatmaps representing the predicted MADRS/HAMD scores following a hypothetical course of TMS treatment to the right dlPFC. Colors represent the predicted change in MADRS/HAMD scores. Y axis represents coil orientation. X axis represents location along the Z-axis of the middle frontal gyrus. The center point of the shaded circle on the heatmaps represents the site and orientation of stimulation predicted to have the maximal reduction in symptoms for each subject. The area of the shaded circle represents the variability (i.e. Euclidean distance [95% confidence interval]) in this optimal site assessed using bootstrapping.

Next we calculated the MADRS change score at the site/orientation combination (5cm anterior to MT at ∼45 degrees eccentricity (30)) typically used for therapeutic rTMS. Consistent with previous literature on remission rates following therapeutic rTMS, our model predicts a reduction in MADRS scores in approximately two-thirds of the patients (n = 59/90) at this site/orientation. Using our approach to optimize site and orientation results in a modest increase in predicted responders (n = 66/90).

### Group-level predictions

To determine whether the model made significant predictions at the group level, we calculated the predicted change in depression symptoms for each site/orientation combination by subtracting the observed pre-TMS MADRS/HAMD from the predicted post-TMS MADRS/HAMD scores. These difference scores represent the predicted change in depression symptoms, given a specific site/orientation of stimulation. We then compared these values to 0 (i.e. no effect of TMS) using a single sample t-test. We corrected for multiple comparisons across sites/orientations the permutation testing procedure described above. As shown in Figure 5 our model predicts a significant decrease in MADRS and HAMD scores in AM subjects with both left and right dlPFC stimulation. In the left hemisphere, the cool spots are centered on BA46 at 30 to 45 degrees orientation. In the right hemisphere, the cool spots are positioned slightly anterior to BA46, and strongest at 120 and 135 degrees orientation. Consistent with the results at the single-subject level, these results suggest that there is a systematic relationship between both site and orientation and predicted symptom change. Importantly, these results are limited to the AM subjects, as the model predicts no significant changes in depression symptoms in the healthy control subjects (See Supplemental Figure 4). In contrast to the data from the dlPFC, the model does not predict any significant changes in depression symptoms for site/orientation combinations centered on the hand region of M1 for either measures (MADRS/HAMD) in AM patients (Supplemental Figure 6) or healthy controls (Supplemental Figure 6).

**Figure 5.**
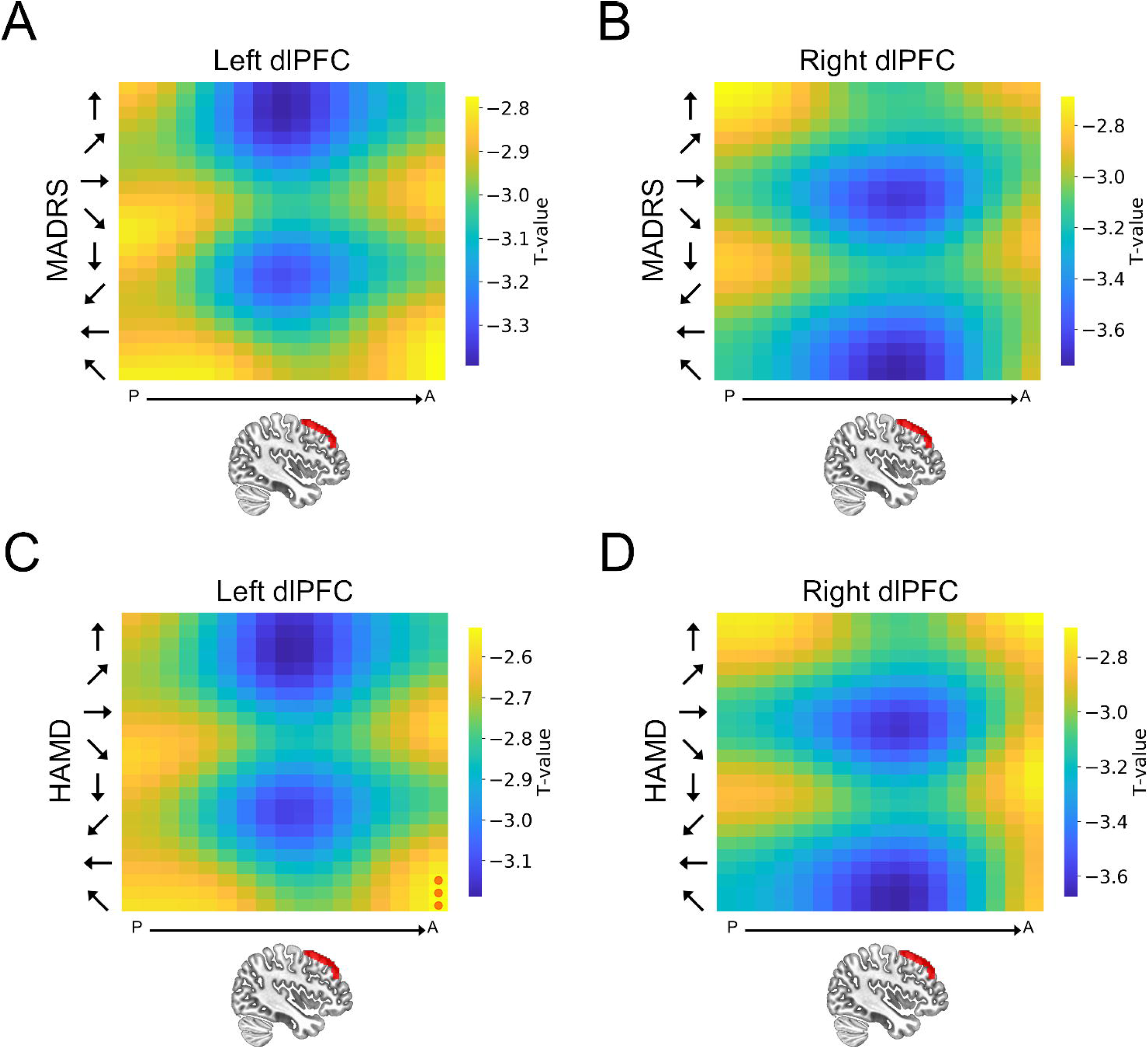
Group-level heatmaps plotting dlPFC predictions for the anxious misery group. **A**) Heatmap representing the predicted MADRS scores following a hypothetical course of TMS treatment to the left dlPFC. **B**) Heatmap representing the predicted MADRS scores following a hypothetical course of TMS treatment to the right dlPFC. **C**) Heatmap representing the predicted HAMD scores following a hypothetical course of TMS treatment to the left dlPFC. **D**) Heatmap representing the predicted HAMD scores following a hypothetical course of TMS treatment to the right dlPFC. Y axis represents coil orientation. Red circles represent sites where the change in MADRS/HAMD scores was not statistically different from 0.

## Discussion

Here we present a proof of concept methodological study where we propose a model that uses functional connectivity, symptom scores, and electric-field modelling to predict symptom change following a hypothetical course of neuromodulatory treatment with TMS. Our model calculates the relationship between symptoms and connectivity, as well as the effect of TMS on connectivity to yield a predicted change score in the symptom, assuming TMS is delivered at the site and orientation specified in the model. We then tested the model systematically across sites and coil orientations in two cortical locations serving as positive (dlPFC) and negative (M1) controls. For the dlPFC, our model suggests that stimulation at this site would lead to a significant reduction in depression symptoms (MADRS/HAMD) in AM patients but not controls, and the magnitude of this potential effect should vary systematically as a function of both site and orientation. Importantly, this observation is specific to the dlPFC, as our model suggests that stimulation at M1 would likely not result in a significant decrease in MADRS or HAMD, regardless of position or orientation. In addition, these observations seem to be reliable at both the single-subject and the group level, suggesting that this approach has sufficient ability to identify subject-specific TMS targets to maximize symptom reduction. The next step for this work is to test the predictions of this model using a variety of therapeutic TMS protocols.

Consistent with previous work implicating the junction of BA9 and BA46 as a potentially optimal site of stimulation for depression symptoms (51–55), our model suggests that stimulation at this site would maximally reduce both MADRS and HAMD scores across subjects compared to the other sites tested (28). Our model adds to the work suggesting that rsFC networks can inform TMS targeting (29), and our results are consistent with the findings that individualized rsFC maps may be most informative (30). Previous research has shown that regions of the dlPFC that are more strongly anticorrelated with sgACC tend to show better clinical efficacy when targeted with therapeutic TMS (51, 56), and the closer the stimulation site is to this optimally anticorrelated dlPFC location, the better the clinical outcome (54, 55, 57). Importantly, active rTMS to the l dlPFC has been shown to reduce anticorrelation between the dlPFC and sgACC (53, 58, 59), and prospective targeting based on individualized dlPFC – sgACC connectivity leads to a greater reduction in symptoms compared to traditional targeting (53, 54). In addition, there is some research to suggest that individualized targeting of BA46 through a parcel guided approach can improve treatment response in patients who have previously failed standard TMS targeted at the dlPFC using the 5-cm rule (28). Based on this work, it could be argued that the current state-of-the-art for individualized TMS targeting for depression is to target sites in the left dlPFC with maximal connectivity with the subgenual cingulate cortex.

However, a review of 33 studies with pre-post resting state shows that although TMS induces robust changes in rsFC, the majority of effects are outside of the stimulated functional network (60). Indeed, the changes in functional connectivity following rTMS treatment for depression can be seen throughout the brain in regions that are important for affective responding (e.g. insula, amygdala, inferior parietal lobule, etc.), suggesting that the net effects on symptoms may be driven by these large scale network connectivity changes (61). Consistent with this idea, treatment response has also been linked to rsFC changes outside of the sgACC (62, 63). Together these results suggest that a whole brain connectivity model may outperform single connection targeting approaches, like the ones described above.

In keeping with these results, our whole brain connectivity approach has several potential benefits over single connection targeting approaches (51–55). First, our model uses e-field modelling to account for the variability in electric field spread due to individual differences in head and cortical anatomy, and uses this information to inform targeting. Second, our approach accounts for other connections in the brain that 1) may be impacted by TMS treatment, and 2) contribute to symptom reduction. Importantly, our model uses a data driven approach to weight these connections by the amount of variability they explain in the depression symptoms. Finally, our approach uses agnostic data-driven estimates between behavioral characteristics and functional connectivity to drive targeting. Therefore, this model can be extended to other symptoms/behavioral characteristics where reliable measures of behavioral markers are available (e.g. addiction, anxiety, etc.).

The main focus of the paper was depressive symptoms, which we captured using scores on the MADRS and HAMD. However, it is important to note that the model described here can be applied to other symptom measures as well. In previous work, we have used various symptom clustering approaches to extract distinct symptom measures from available clinical data (64), resulting in several distinct symptom clusters. Although out of the scope of the current work, future studies should determine the degree to which these other distinct symptom clusters could potentially be used to direct TMS targeting. Indeed, defining biotypes for depression and anxiety starting with neural circuit dysfunction (65), may yield additional symptom-specific targets. Accordingly, the model presented here could be used not only to individualize targeting at the single subject level, but it could potentially be extended to individualize targeting at the single session level, where each TMS session in a course of treatment is targeted at a specific symptom or set of symptoms, thereby maximizing the therapeutic effect.

### Strengths and Limitations

The strength of the work described here stems from the generality of its applicability. The model described in this work is both site and symptom independent. Therefore, the methods described here can be used to optimize TMS targeting for a variety of disorders shown to be effectively treated with TMS (66). This approach could be used prospectively to screen for potential TMS effects within new disorders or for novel symptom dimensions. The model is also able to make predictions at the single-subject level regarding the optimal stimulation site for a given symptom, allowing for single-subject optimization of TMS targeting. In all cases, the predictions of the model can be directly tested by applying TMS according to the model predictions. Another strength is that our results are consistent for MADRS and HAMD scores, suggesting that our model is capable of identifying stimulation sites to target depressive symptoms. However, it should be noted MADRS and HAMD scores are highly correlated across subjects (AM, r = 0.857; Control, r = 0.664), which means that the similarity in results across the two measures is likely driven by this shared variability. Accordingly, it is important not to characterize the results from these two scores as independent results, but rather different characterizations of the same underlying relationship between depression and rsFC.

The primary weakness of this study is that it is a proof of concept study, rather than a confirmatory study, meaning we do not actually administer TMS and measure the pre/post symptom changes in the current work. We understand validation of the current model is the necessary next step. Publication of the model in its initial state will allow multiple labs to test the predictions of the model, allowing for a more rapid validation/refinement of the work. To facilitate this process, we will make the code for our model publicly available.

Another potential weakness of the current work is that we used Pearson’s correlations, a non-directional measure, to model functional connectivity. However, it is likely that TMS affects upstream and downstream connections differentially (67). Accordingly, it may be more appropriate to model the functional connections using effective connectivity, and differentially weight the connections based on whether they are upstream or downstream of the stimulation site. Of course this becomes more complicated in our model, given that the stimulation “site” is modelled as a thresholded map of the current induced across the whole brain, given a TMS coil position and orientation. Accordingly, to factor in differential effects for upstream and downstream connections, it would be necessary to generate an effective connectivity map of the entire brain. While techniques like dynamic causal modelling (DCM) can be used to generate realistic effective connectivity models with a limited number of sites and connections (68–70), it remains to be seen whether this approach can be reliably used to generate whole brain effective connectivity models for use at the single-subject level (71).

Another limitation is that the model assumes that changes in connectivity tend to scale linearly with the magnitude of the electric field induced, given a particular site/orientation of stimulation. This mathematical assumption likely holds for little if any *in vivo* applications of TMS. Indeed, it is well known that the effects of TMS on outcome variables like cortical excitability (72, 73), functional connectivity (53, 59), and behavior (74, 75) are dependent on the stimulation parameters used (60, 76). Indeed, amplitude (77), frequency (78), pattern (79), number of sessions (6), and intervals (79) between sessions are all known to be important determinants of observed TMS effects. We understand that the linear assumption put forth in our model is a massive oversimplification, but we see this as a starting point. Accordingly, we have included a proportionality constant in our model to accommodate parameter-dependent scaling in connectivity changes. Our goal is to state the initial model as parsimoniously as possible, to allow the model to be tested with a variety of experimental and clinical TMS protocols. As more information becomes known about the relationship between rTMS parameters and connectivity changes, researchers will be able to adjust the proportionality constant of the model to suit their protocol.

### Conclusions

Individualized targeting is one key to maximizing the efficacy of clinical rTMS, and mapping TMS-induced changes in rsFC to changes in clinical symptoms is critically important for this goal. The model proposed is one potential approach to accomplish this goal. In this initial work, we show that the model makes predictions at both positive and negative control sites that are consistent with observed symptom changes following therapeutic applications of rTMS for depression (51–55). The primary strength of this model is the generality of our approach, as the model proposed is both site and symptom independent. Our hope is that this work will be applied more generally to optimize TMS targeting across psychiatric disorders.

## Author contributions

**NLB:** Designed Study, Conducted analysis, wrote manuscript, provided supervision, edited manuscript. **JCB:** Conducted statistical analysis; edited manuscript. **DS:** Assisted with data collection and preprocessing. **WM:** Assisted with data collection and preprocessing. **ZD:** Assisted with statistical analysis and manuscript preparation. **TG:** Assisted with data collection and preprocessing,. **MT:** Assisted with data collection and preprocessing, assisted with manuscript preparation. **NS:** Assisted with data collection and preprocessing. **MJ:** Assisted with data collection and preprocessing. **DJO:** Assisted with study design; edited manuscript. **YIS:** Assisted with study design, wrote manuscript; provided supervision; edited manuscript.

## Acknowledgments

This study utilized the high-performance computational capabilities of the Cubic computing cluster at the University of Pennsylvania. (https://www.med.upenn.edu/cbica/cubic.html). The authors would like to thank Maria Prociuk for her expertise and assistance in submitting the manuscript. We would also like to thank the participants for their time and effort.

## Ethical Standards

The authors assert that all procedures contributing to this work comply with the ethical standards of the relevant national and institutional committees on human experimentation and with the Helsinki Declaration of 1975, as revised in 2008.

