## Supplement for "Proof of concept study to develop a novel connectivity-based electric-field modelling approach for individualized targeting of transcranial magnetic stimulation treatment"

**Running title:** Novel modelling approach for individualized TMS targeting

**Authors:**	Nicholas L Balderston^1^, Joanne C Beer^2^, Darsol Seok^1^, Walid Makhoul^1^, Zhi-De Deng^3^, Tommaso Girelli^1^, Marta Teferi^1^, Nathan Smyk^1^, Marc Jaskir^1^, Desmond J Oathes^1^, & Yvette I Sheline^1^

1. Center for Neuromodulation in Depression and Stress
   Department of Psychiatry
   University of Pennsylvania
   Philadelphia, PA, USA
2. Penn Statistics in Imaging and Visualization Center

Department of Biostatistics, Epidemiology & Informatics

University of Pennsylvania

Philadelphia, PA, USA

3. Noninvasive Neuromodulation Unit

National Institute of Mental Health

Bethesda, MD, USA

**Corresponding Author:**

Nicholas Balderston

Center for Neuromodulation in Depression and Stress

3700 Hamilton Walk, Richards D302

Philadelphia, PA, 19104

**Keywords:** Transcranial Magnetic Stimulation, resting state, functional connectivity, electric field modelling, depression, anxiety, anxious misery

**Appendix: Equations**

Connectivity based model: PCA regression

Let $\text{Y}^{mood}$ be the $n \times1$ vector of behavioral symptom scores (e.g., MADRS scores) for $n$ participants. We assume the behavioral symptom scores can be predicted using a linear combination of connectivity measures, according to the model

$$\boldsymbol{Y}^{mood}=\boldsymbol{X\beta+\epsilon} (Eq. A1)$$

where $\boldsymbol{X}$ is a $n \times p$ matrix such that each row consists of one participant’s concatenated unique connectivity values from the Z-transformed Pearson correlation functional connectivity matrix, i.e., the upper or lower triangular portion of the $N\times N$ connectivity matrix ($p=N\left( N-1 \right)/2$), for the Gordon atlas, $N=333$); $\boldsymbol{\beta}$ is a $p \times1$ vector of coefficients, and $\boldsymbol{\epsilon}$ is a $n \times1$ vector of error terms. We assume that $\boldsymbol{Y}^{mood}$ and columns of $\boldsymbol{X}$ have been centered so as to have zero empirical means.

We perform principal components analysis (PCA) on the predictor matrix $\boldsymbol{X}$. Let $\boldsymbol{X=U\Delta}\boldsymbol{V}^{\boldsymbol{T}}$ denote the singular value decomposition of $\boldsymbol{X}$, where $\boldsymbol{\Delta}_{p\times p}=\text{diag}\left( \delta_{1},\ldots,\delta_{p} \right)$ is a diagonal matrix of non-negative singular values, and columns of $\boldsymbol{U}_{n\times p}$ and $\boldsymbol{V}_{p\times p}$ are orthonormal sets of vectors, i.e., the left and right singular vectors of $\boldsymbol{X}$. The spectral decomposition of $\boldsymbol{X}$ is given by $\boldsymbol{V\Lambda}\boldsymbol{V}^{\boldsymbol{T}}$, where $\boldsymbol{\Lambda}_{p\times p}=\text{diag}\left( \lambda_{1},\ldots,\lambda_{p} \right)=\text{diag}\left( \delta_{1}^{2},\ldots,\delta_{p}^{2} \right)=\boldsymbol{\Delta}^{2}$ is a diagonal matrix of the non-negative eigenvalues of $\boldsymbol{X}^{\boldsymbol{T}}\boldsymbol{X}$ and columns of $\boldsymbol{V}_{p\times p}=\left[ \boldsymbol{v}_{\boldsymbol{1}}\boldsymbol{,\ldots,}\boldsymbol{v}_{\boldsymbol{p}} \right]$ are the corresponding eigenvectors. The j^th^ principal component and j^th^ principal component direction (i.e., PCA loading) corresponding to the j^th^ largest eigenvector are $\boldsymbol{X}\boldsymbol{v}_{\boldsymbol{j}}$ and $\boldsymbol{v}_{\boldsymbol{j}}$, respectively, for each $j\in\left\{ 1, \ldots, p \right\}$.

For PCA regression (PCR), we use the first $k\leq n$ components as predictors. Let $\boldsymbol{V}_{k}$ represent the $p \times k$ matrix comprising the first $k$ columns of $\boldsymbol{V}$, and $\boldsymbol{W}_{k}=\boldsymbol{X}\boldsymbol{V}_{k}$ be the $n \times k$ matrix with the first $k$ principal components as columns. We estimate regression coefficients $\boldsymbol{\gamma}_{k}$ in $\boldsymbol{Y}^{mood}=\boldsymbol{W}_{k}\boldsymbol{\gamma}_{k}\boldsymbol{+}\boldsymbol{\epsilon}_{k} (Eq. A2)$ by ordinary least squares, i.e., $\boldsymbol{\gamma}_{k}=\left( \boldsymbol{W}_{k}^{T}\boldsymbol{W}_{k} \right)^{-1}\boldsymbol{W}_{k}^{T}\boldsymbol{Y}^{mood}\in\mathbb{R}^{k}$. Then the PCR estimator of $\boldsymbol{\beta}$ based on the first $k$ principal components is given by ${\hat{\boldsymbol{\beta}}}_{k}=\boldsymbol{V}_{k}{\hat{\boldsymbol{\gamma}}}_{k}\in\mathbb{R}^{p} (Eq. A3)$.

E-field augmented model

Once ${\hat{\boldsymbol{\beta}}}_{k}$ is obtained from the connectivity based model, the e-field augmented model will be used to identify the optimal coil orientation and stimulation sites to generate the greatest reduction in behavioral symptom score for each participant. Let $\Delta\boldsymbol{Y}^{mood}$ be the $n \times1$ vector of changes in score between the pre- and post-treatment assessments for $n$ participants. We model $\Delta\boldsymbol{Y}^{mood}$ as follows:

$$\Delta\boldsymbol{Y}^{mood}=\boldsymbol{Y}_{post}^{mood}-\boldsymbol{Y}_{pre}^{mood}=\left( \boldsymbol{X}_{post}\boldsymbol{-}\boldsymbol{X}_{pre} \right)\boldsymbol{\beta+}\boldsymbol{\epsilon}^{\boldsymbol{'}} (Eq. A4),$$

where $\boldsymbol{\epsilon'}=\boldsymbol{\epsilon}_{post}-\boldsymbol{\epsilon}_{pre}$.

For each site and orientation $l$, let $\boldsymbol{E}_{l}$ be a $n \times p$ matrix, where rows correspond to participants, columns correspond to pairs of regions in the Gordon atlas, and entries equal the average e-field model values at each pair of regions for the given participant, site, and orientation. We assume that $\boldsymbol{X}_{post}^{l}\boldsymbol{=}C\cdot\boldsymbol{E}_{l}\circ\boldsymbol{X}_{pre}$, where $C$ is a positive proportionality constant, and $\boldsymbol{\circ}$ denotes the Hadamard product. Thus, for a given stimulation site and coil orientation combination $l$,

$$\Delta\boldsymbol{Y}^{mood}=\left( C\cdot\boldsymbol{E}_{l}\circ\boldsymbol{X}_{pre}\boldsymbol{-}\boldsymbol{X}_{pre} \right)\boldsymbol{\beta+}\boldsymbol{\epsilon}^{\boldsymbol{'}} (Eq. A5)\boldsymbol{.}$$

With these assumptions, and substituting the ${\hat{\boldsymbol{\beta}}}_{k}$ estimated form the PCR described above, it is possible to compare relative differences in the predicted change in behavioral symptom score across sites and orientations $l$.

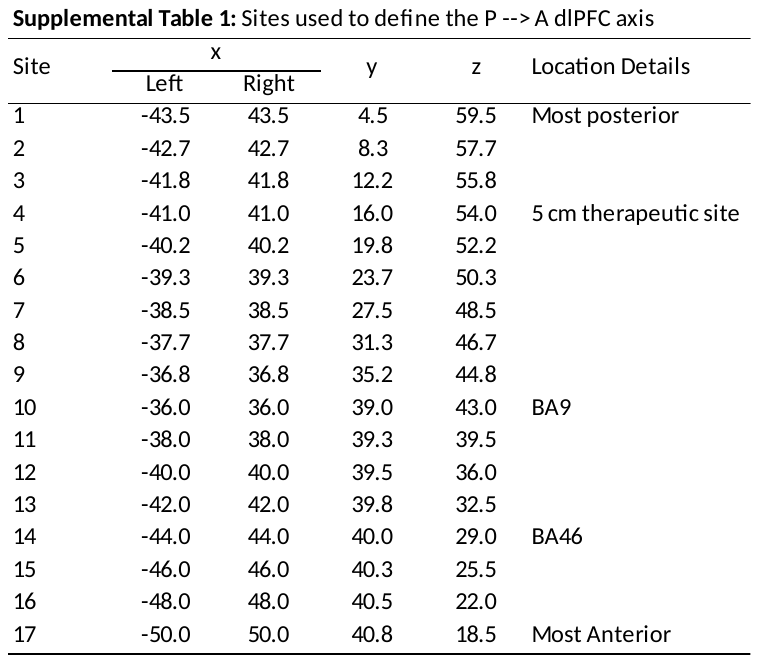

**Note: Coordinates in MNI space.**

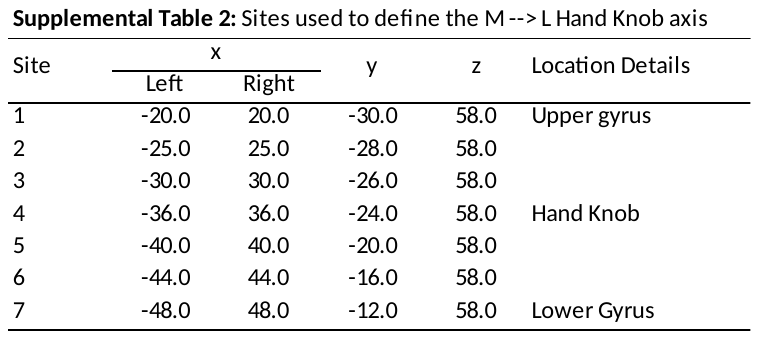

**Note: Coordinates in MNI space.**

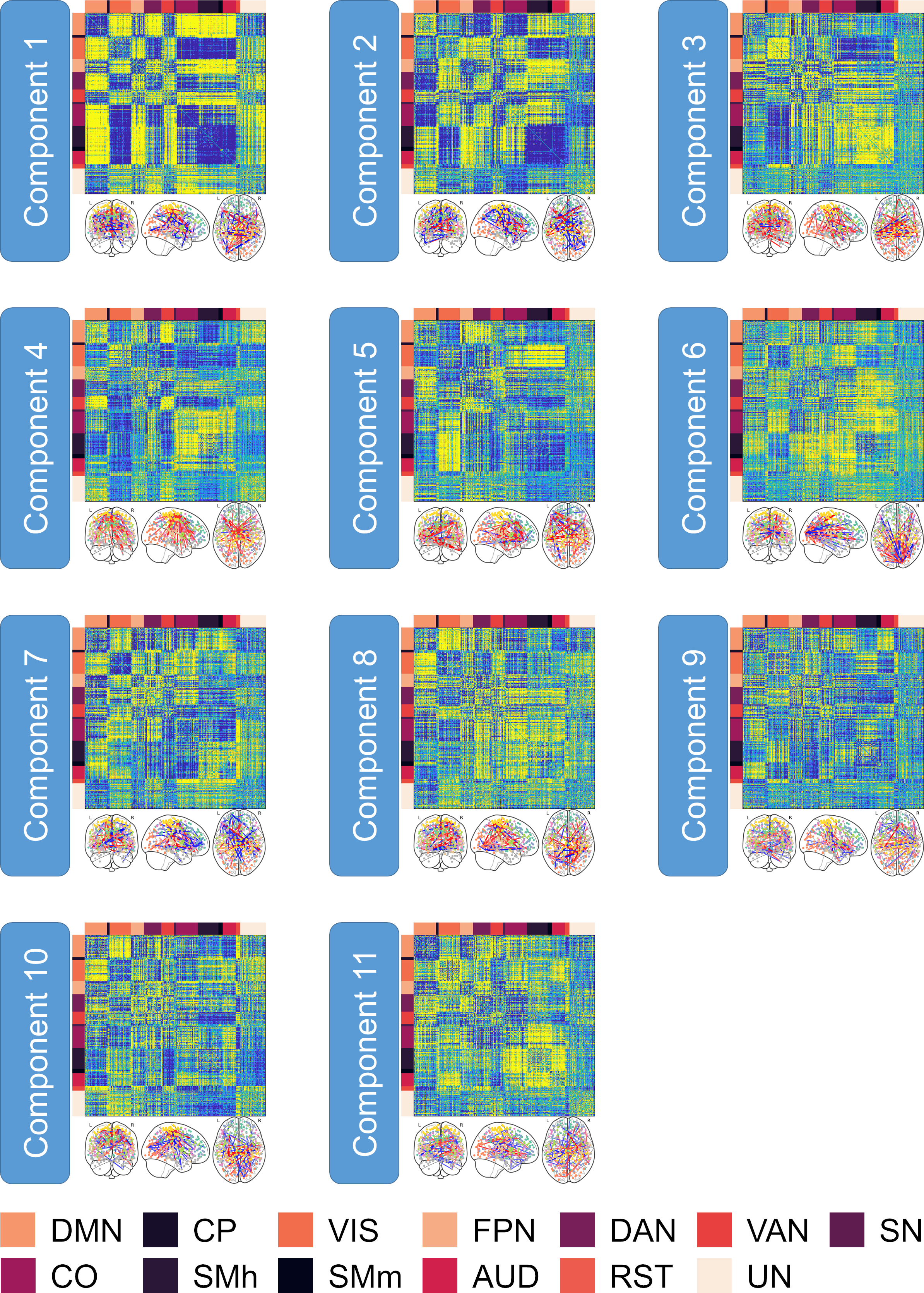

**Supplemental Figure 1. Item coefficients for the components selected from the Anxious Misery Principle Components Analysis.** Matrices represent the loading of each pairwise connection onto the principle components, sorted by network according to the key at the bottom of the figure. Connectome graphs for each of the principle components show the top 0.1% of connections according to the item weightings. **Network Color key**: **DMN** = Default Mode Network; **CP** = CinguloParietal; **VIS** = Visual; **FPN** = FrontoParietal Network; **DAN** = Dorsal Attention Network; **VAN** = Ventral Attention Network; **SN** = Salience Network; **CO** = CinguloOpercular; **SMh** = SomatoMotor (hand); **SMm** = SomatoMotor (mouth); **AUD** = Auditory; **RST** = RetrosplenialTemporal; **UN** = Unassigned nodes.

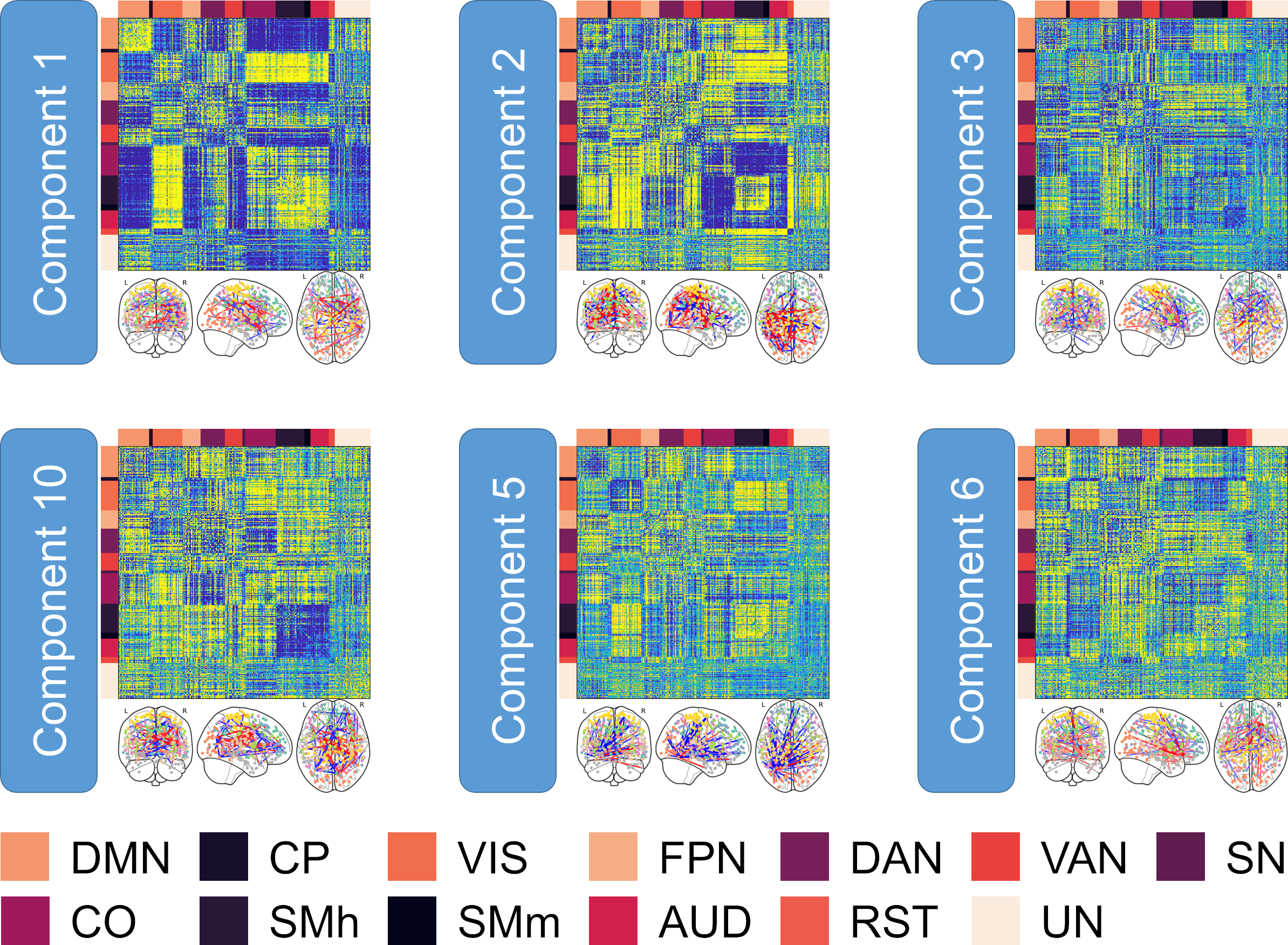

**Supplemental Figure 2. Item coefficients for the components selected from the Healthy Control Principle Components Analysis.** Matrices represent the loading of each pairwise connection onto the principle components, sorted by network according to the key at the bottom of the figure. Connectome graphs for each of the principle components show the top 0.1% of connections according to the item weightings. **Network Color key**: **DMN** = Default Mode Network; **CP** = CinguloParietal; **VIS** = Visual; **FPN** = FrontoParietal Network; **DAN** = Dorsal Attention Network; **VAN** = Ventral Attention Network; **SN** = Salience Network; **CO** = CinguloOpercular; **SMh** = SomatoMotor (hand); **SMm** = SomatoMotor (mouth); **AUD** = Auditory; **RST** = RetrosplenialTemporal; **UN** = Unassigned nodes.

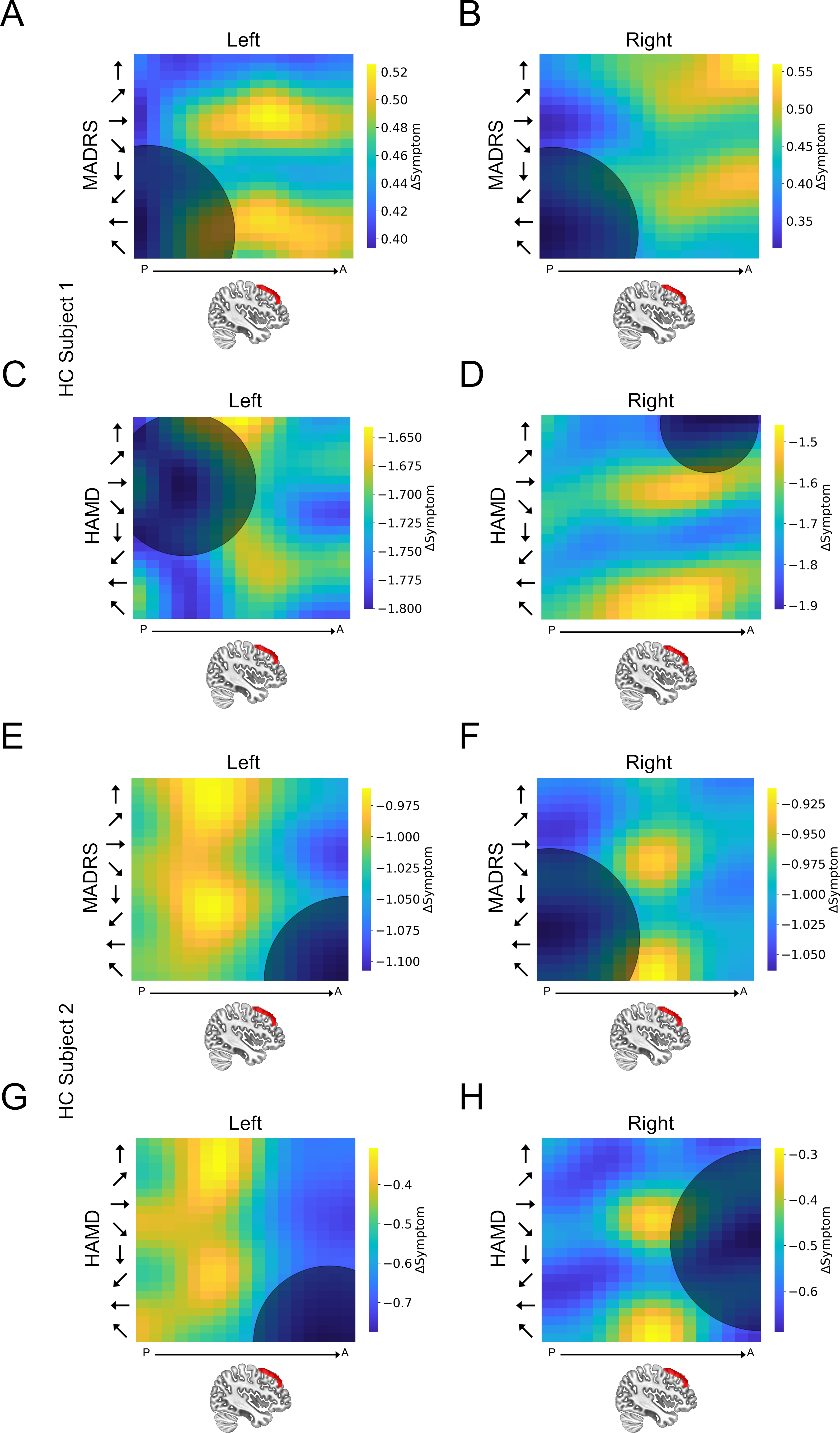

**Supplemental Figure 3. Individual subject heatmaps plotting dlPFC predictions for the healthy control group.** (**A, C, E, G**) Heatmaps representing the predicted MADRS and HAMD scores following a hypothetical course of TMS treatment to the left dlPFC. (**B, D, F, H**) Heatmaps representing the predicted MADRS/HAMD scores following a hypothetical course of TMS treatment to the right dlPFC. Colors represent the predicted change in MADRS/HAMD scores. Y axis represents coil orientation. X axis represents location along the Z-axis of the middle frontal gyrus. The center point of the shaded circle on the heatmaps represents the site and orientation of stimulation predicted to have the maximal reduction in symptoms for each subject. The area of the shaded circle represents the variability (i.e. Euclidean distance [95% confidence interval]) in this optimal site assessed using bootstrapping.

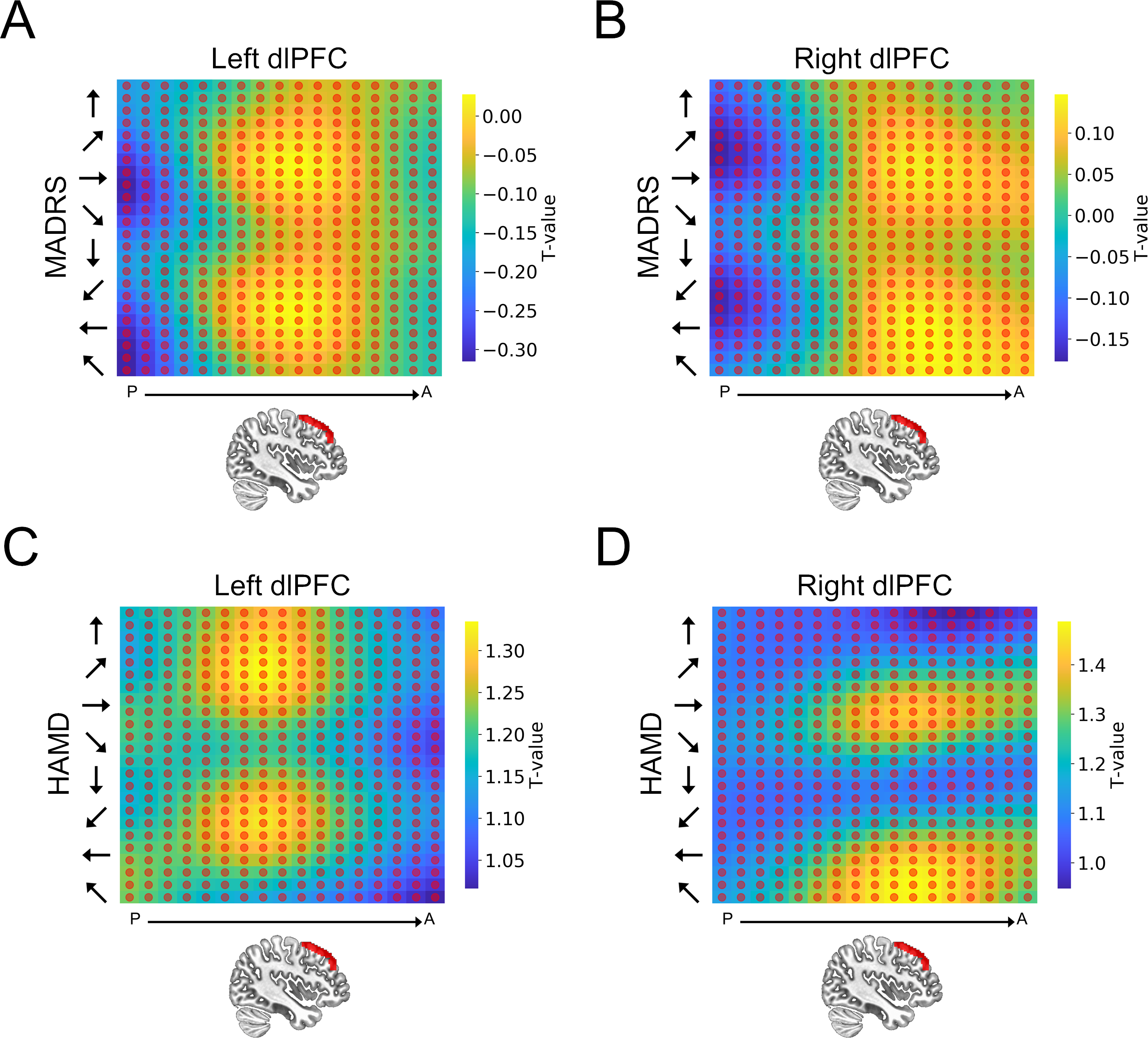

**Supplemental Figure 4. Group-level heatmaps plotting dlPFC predictions for the healthy control group. A**) Heatmap representing the predicted MADRS scores following a hypothetical course of TMS treatment to the left dlPFC. **B**) Heatmap representing the predicted MADRS scores following a hypothetical course of TMS treatment to the right dlPFC. **C**) Heatmap representing the predicted HAMD scores following a hypothetical course of TMS treatment to the left dlPFC. **D**) Heatmap representing the predicted HAMD scores following a hypothetical course of TMS treatment to the right dlPFC. Y axis represents coil orientation. X axis represents location along the Z-axis of the middle frontal gyrus. Red circles represent sites where the change in MADRS/HAMD scores was not statistically different from 0.

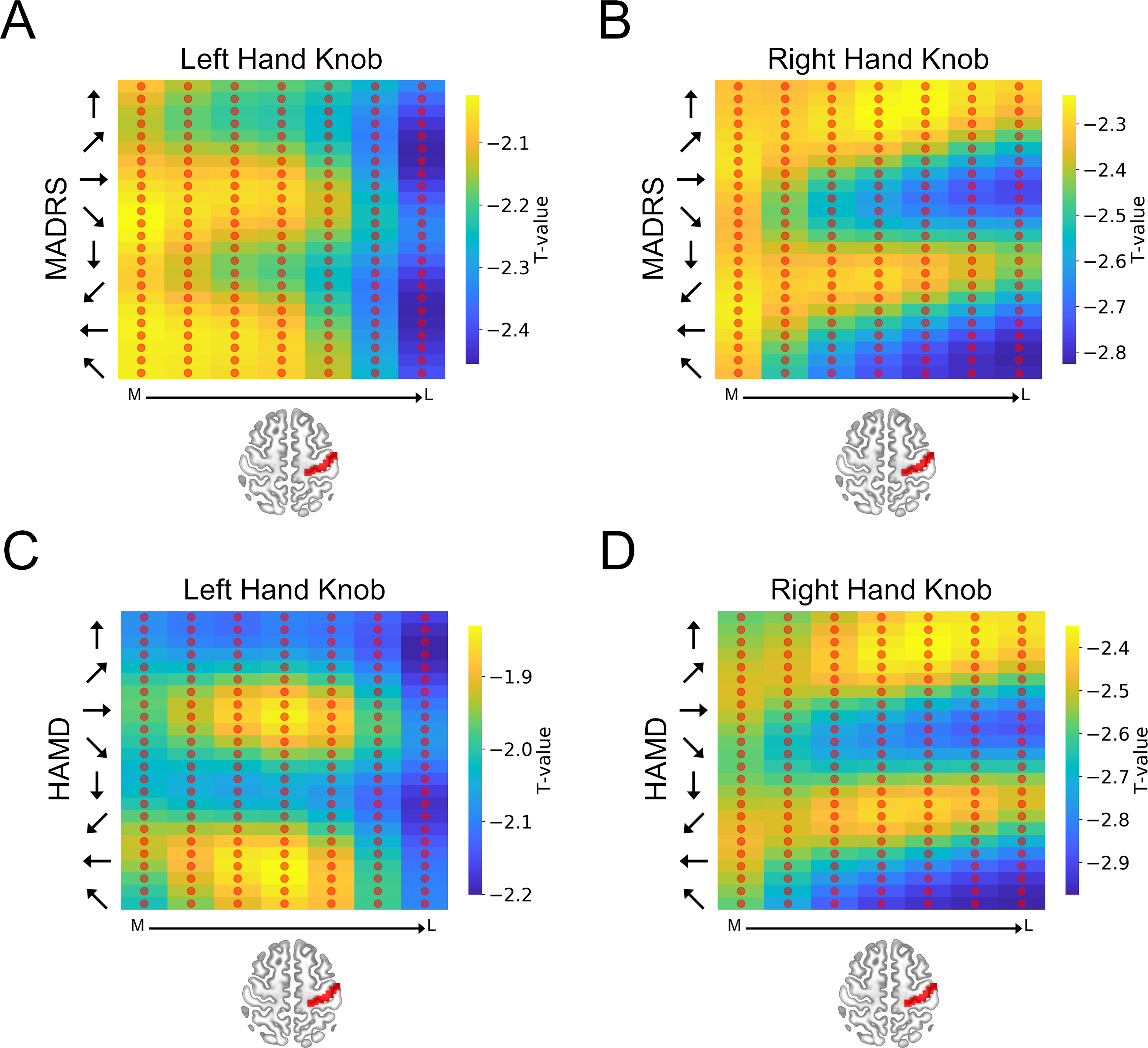

**Supplemental Figure 5. Group-level heatmaps plotting hand knob predictions for the anxious misery group. A**) Heatmap representing the predicted MADRS scores following a hypothetical course of TMS treatment to the left hand knob. **B**) Heatmap representing the predicted MADRS scores following a hypothetical course of TMS treatment to the right hand knob. **C**) Heatmap representing the predicted HAMD scores following a hypothetical course of TMS treatment to the left hand knob. **D**) Heatmap representing the predicted HAMD scores following a hypothetical course of TMS treatment to the right hand knob. Y axis represents coil orientation. X axis represents location along the medial to lateral axis of the primary motor cortex. Red circles represent sites where the change in MADRS/HAMD scores was not statistically different from 0.

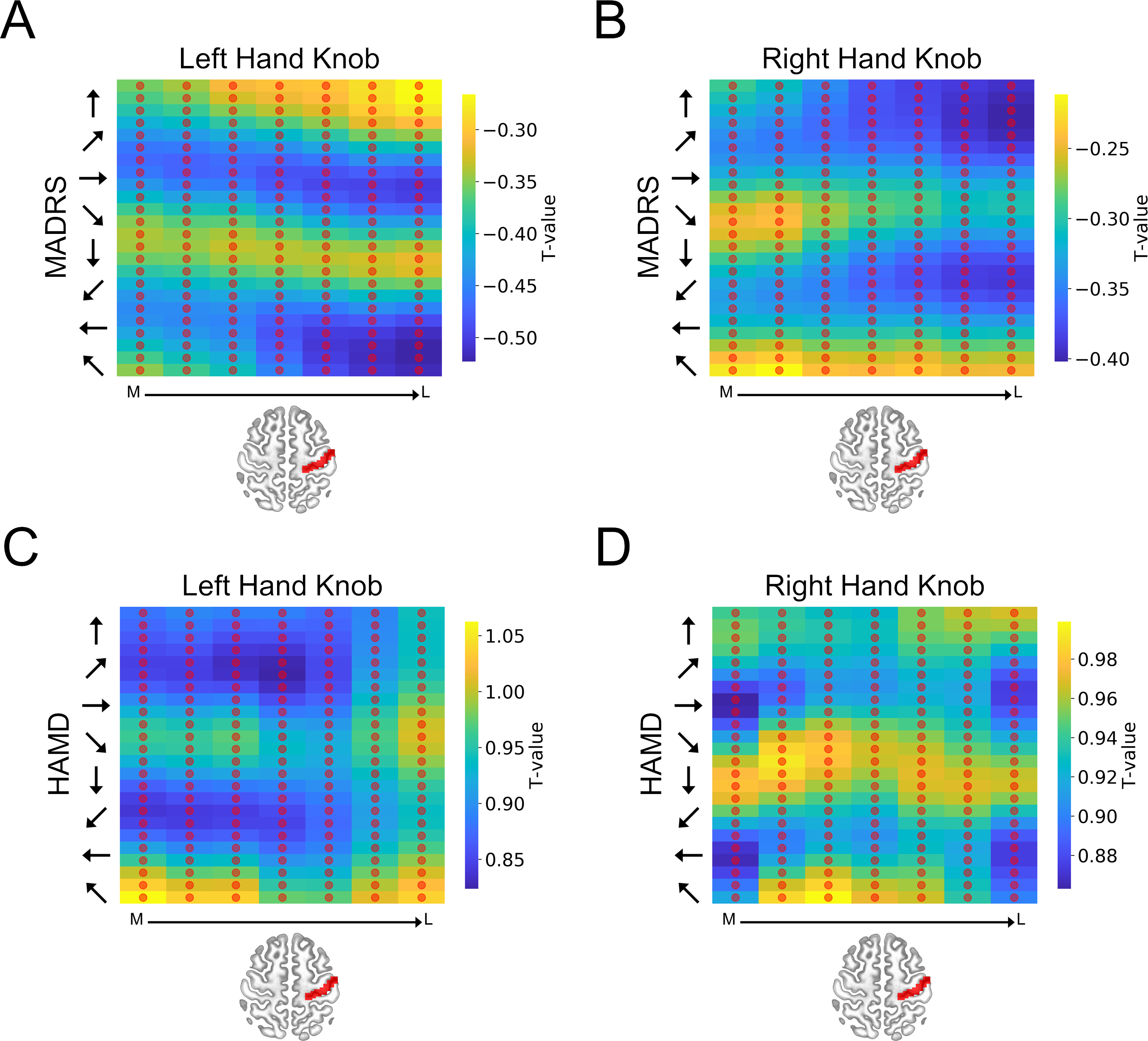

**Supplemental Figure 6. Group-level heatmaps plotting hand knob predictions for the healthy control group. A**) Heatmap representing the predicted MADRS scores following a hypothetical course of TMS treatment to the left hand knob. **B**) Heatmap representing the predicted MADRS scores following a hypothetical course of TMS treatment to the right hand knob. **C**) Heatmap representing the predicted HAMD scores following a hypothetical course of TMS treatment to the left hand knob. **D**) Heatmap representing the predicted HAMD scores following a hypothetical course of TMS treatment to the right hand knob. Y axis represents coil orientation. X axis represents location along the medial to lateral axis of the primary motor cortex. Red circles represent sites where the change in MADRS/HAMD scores was not statistically different from 0.
